# Neural envelope tracking predicts speech intelligibility and hearing aid benefit in children with hearing loss

**DOI:** 10.1101/2023.07.03.547477

**Authors:** Tilde Van Hirtum, Ben Somers, Benjamin Dieudonné, Eline Verschueren, Jan Wouters, Tom Francart

## Abstract

Early assessment of hearing aid benefit is crucial, as the extent to which hearing aids provide audible speech information predicts speech and language outcomes. A growing body of research has proposed neural envelope tracking as an objective measure of speech, particularly for individuals unable to provide reliable behavioral feedback. However, its potential for evaluating speech intelligibility and hearing aid benefit in hearing-impaired children remains unexplored. This study examined neural envelope tracking in hearing-impaired children through two separate experiments. EEG data was recorded while children listened to age-appropriate stories (experiment 1) or an animated movie (experiment 2) under aided and unaided conditions (using personal hearing aids) at multiple stimulus intensities. Results in the delta band demonstrated that neural tracking increased with increasing stimulus intensity, but only in the unaided condition. In the aided condition, neural tracking remained stable across a wide range of intensities, as long as speech intelligibility was maintained. This suggests that acoustic degradation of the speech signal does not necessarily impact neural tracking. Additionally, the use of personal hearing aids significantly enhanced neural envelope tracking, particularly in challenging speech conditions (which would be inaudible when unaided). Furthermore, neural envelope tracking strongly correlated with behaviorally measured speech intelligibility. Altogether, these findings indicate that neural envelope tracking could be a valuable tool for predicting speech intelligibility benefits derived from personal hearing aids in hearing-impaired children. Incorporating narrated stories or engaging movies expands the accessibility of these methods even in clinical settings, offering new avenues for using objective speech measures to guide pediatric audiology decision-making.

## Introduction

Permanent childhood hearing loss affects 1 to 2 per 1000 newborns globally (Lieu et al., 2020). When hearing loss goes unidentified, children are at risk of experiencing delays in speech-language development, socio-emotional challenges, difficulties with cognitive abilities, and academic underachievement (World Health Organization, 2016). The implementation of universal newborn hearing screening has significantly lowered the average age of identification. Early detection facilitates early intervention, which allows for timely fitting of hearing aids to improve access to spoken language within a benchmark of no later than 3 to 6 months (The Joint Committee on Infant Hearing, 2019).

The degree to which hearing aids provide audible speech information is a key predictor of speech and language outcomes (Stiles et al., 2012; Tomblin et al., 2014; Walker et al., 2019). Furthermore, the benefits of aided hearing contribute to the resilience of children with hearing loss, helping them achieve academic performance comparable to their normal-hearing peers (Tomblin et al., 2020). Therefore, early assessment of hearing aid benefit is essential. While the evaluation of hearing aid fitting can be established with reasonable certainty in individuals who are able to respond reliably to conventional behavioral testing methods, it becomes more challenging or even impossible in infants, young children or children with special needs.

The recognition of the clinical necessity for assessing the benefit of hearing aids has led to the development of objective electroencephalogram (EEG)-based measures, which do not require active participation from the listener. Previous research indicates that (aided) event-related potentials (ERPs) such as cortical auditory evoked potentials (CAEPs) can be a valuable tool for evaluating the functionality of hearing aids (Punch et al., 2016). Unlike auditory brainstem responses (ABRs), which rely on brief tones or clicks, CAEPs can be elicited using speech stimuli such as short phonemes or syllables. For instance, Van Dun et al. (2012) recorded CAEPs in response to three brief frequency-distinct speech sounds /m/, /g/, and /t/ and demonstrated that the presence (or absence) of CAEP responses is related to the audibility of these speech sounds for hearing-impaired infants. Moreover, multiple studies have reported an increase in CAEP amplitudes and a greater number of detected responses when comparing unaided and aided testing (Chang et al., 2012; Van Dun et al., 2016). Differences between unaided and aided conditions were more likely to occur near threshold rather than at suprathreshold levels (Korczak et al., 2005), thus suggesting that amplification indeed enhances CAEP amplitudes, particularly when unaided speech sounds are barely audible or not audible at all.

In addition to CAEPs, the practical feasibility of using speech-evoked envelope following responses (EFRs) to assess hearing aid benefit has been evaluated. Speech-evoked EFRs are steady-state responses that reflect neural responses phase-locked to the envelope periodicity in phonemes. Naturally spoken as well as synthetic vowels can elicit EFRs at the fundamental frequency of the voice (Aiken and Picton, 2006). In contrast to CAEPs that use brief phonemes as the stimulus along with interstimulus intervals of 1 s (e.g. Chang et al., 2012), the main advantage of the speech-evoked EFR paradigm is the use of a longer, continuous stimulus. This is preferable to avoid undesired interactions with nonlinear hearing aid function (Scollie and Seewald, 2002; Easwar et al., 2012). The amplitude and detectability of EFRs was found to increase with the use of hearing aids for both hearing-impaired adults (Anderson and Kraus, 2013; Easwar et al., 2015) and children (Easwar et al., 2023), similarly to the findings observed in CAEP studies.

Taken together, CAEP and EFR studies with adults and children illustrate that detection of a response can predict audibility of speech sounds. Moreover, a clear relation with behavioral thresholds as well as tone-burst ABR results has been demonstrated (e.g. Van Dun et al., 2012; Chang et al., 2012; Baydan et al., 2019). As a result, an improvement in response detectability while using hearing aids could potentially serve as a reliable predictor of the benefits derived from hearing aid usage. However, it remains unclear whether these AEP are directly related with measures of speech intelligibility, especially in hearing-impaired listeners. In addition, multiple repetitions of identical short stimuli are required to evoke cortical responses, which do not reflect ecologically relevant stimuli that are encountered in daily life (Alexandrou et al., 2020).

A better evaluation of speech intelligibility and the functionality of the hearing aid in daily life could be achieved by assessing neural speech tracking. This approach allows the use of running speech, such as audiobooks or podcasts (for a review, see Brodbeck and Simon, 2020), by analyzing the mathematical relation between the speech dynamics and the corresponding neural responses in a listener’s brain (e.g. Ding and Simon, 2012b; Gross et al., 2013; Ding et al., 2016). Given that the speech envelope carries crucial acoustic information required for spoken language comprehension (Shannon et al., 1995), a variety of studies have investigated whether neural envelope tracking can be used as a measure of speech intelligibility. Accordingly, a growing body of literature shows that neural activity tracks the temporal fluctuations of the speech envelope during speech perception in adults (e.g. Ding and Simon, 2012a, 2013; O’Sullivan et al., 2015; Vanthornhout et al., 2018; Verschueren et al., 2021), as well as children (Vander Ghinst et al., 2019; Ríos-López et al., 2020; Van Hirtum et al., 2023) and infants (e.g. Kalashnikova et al., 2018; Attaheri et al., 2022. More importantly, multiple studies have shown a positive correlation of neural envelope tracking and behavioral measures of speech intelligibility in normal hearing listeners, particularly in the delta frequency range (0.5-4 Hz) (e.g. Ding et al., 2014; Iotzov and Parra, 2019; Vanthornhout et al., 2018; Lesenfants et al., 2019; Verschueren et al., 2021; Van Hirtum et al., 2023).

However, only a limited number of studies have investigated the effect of hearing loss and/or the use of a hearing aid on neural tracking of the speech envelope, and so far only with (older) adults. These studies confirm the positive correlation of neural envelope tracking with speech intelligibility (Verschueren et al., 2019; Decruy et al., 2020; Paul et al., 2020), although it is worth noting that neural envelope tracking in hearing-impaired listeners is generally higher compared to normal-hearing peers (Decruy et al., 2020; Mirkovic et al., 2019; Fuglsang et al., 2020, although see Petersen et al., 2017; Presacco et al., 2019). To minimize the effect of differences in audibility, most of the above-mentioned studies amplified the presented speech stimuli in a frequency-dependent way based on the audiogram (e.g. Decruy et al., 2020; Fuglsang et al., 2020). This approach simulates the situation of aided listening (e.g., with a linear hearing aid). Research on the potential effect of (personal) hearing aids on neural tracking of speech is scarce, although better audibility is expected to result in a enhanced neural envelope tracking. Only one study to our knowledge has studied neural tracking of the speech envelope with and without hearing aids in hearing-impaired adults. Vanheusden et al. (2020) found that neural tracking to speech at an audible level (i.e. 70 dB A) is not affected by hearing aids in older adults with mild-to-moderate hearing loss. Their results showed no significant differences between unaided and aided conditions in the number of responses detected, nor in the strength of neural tracking. Therefore, the authors concluded that hearing aids do not significantly alter neural tracking under fully audible conditions. In this regard, multiple studies so far provided evidence that neural tracking in the delta band remains stable, as long as the acoustical degradation of the speech signal (i.e. lower stimulus intensity or more background noise) does not reflect significant changes in speech intelligibility (Ding and Simon, 2013; Verschueren et al., 2021; Van Hirtum et al., 2023). Taken together, these findings indicate that differences in neural tracking are not purely driven by amplification, showing the potential of using neural tracking to evaluate hearing function as well as hearing aid benefit.

To date, the same approach has not been used with hearing-impaired children. On the contrary, the majority of electrophysiological studies investigating the effect of hearing loss and/or hearing aids have involved older adults. In the present study, we therefore investigate the effect on neural envelope tracking in children with permanent hearing loss of (i) stimulus intensity and (ii) the use of personal hearing aids, and (iii) whether neural envelope tracking reflects speech intelligibility in these children. We investigate neural tracking by combining both linear decoding (backward-modelling) and encoding (forward-modelling) models, providing complementary information about neural tracking (for a review see Gillis et al., 2022b. The backward modelling approach uses a linear model to reconstruct the acoustic representation, here the envelope, based on the measured EEG responses. This results in a reconstruction accuracy, which indicates how well the envelope is reconstructed from the EEG responses. With the forward modelling approach, a linear model predicts the EEG responses to speech and yields a temporal response function (TRF). Such a TRF can be used to study the spatio-temporal dynamics of the response. The feasibility of this approach in young children was shown recently by Van Hirtum et al. (2023), and this is the first study to use this method in hearing-impaired children.

Here, we conducted two separate experiments. In the first experiment, we selected narrated stories accompanied by story illustrations as the stimulus, following a similar protocol and methods as Van Hirtum et al. (2023). The stories were presented in quiet at sound intensities well above (i.e. audible) and below (inaudible) hearing thresholds, to investigate whether the application of hearing aids affects neural tracking of speech. Although running speech from narrated stories is often used, it still differs from everyday speech and natural dialogues. Therefore, in the second experiment, we assessed the feasibility of using an animated movie as the stimulus. With this approach, we aim to capture different aspects and contexts of real-life speech usage. For both experiments, we have three specific hypotheses: First, we hypothesize that as stimulus intensity increases, neural envelope tracking increases, given that the changes in stimulus intensity reflect changes in speech intelligibility (Ding and Simon, 2013; Verschueren et al., 2021; Van Hirtum et al., 2023). Next, we hypothesize that amplification by means of personal hearing aids enhances neural tracking. Previous research has shown higher neural responses when comparing unaided and aided testing when unaided speech sounds were barely audible (e.g. Van Dun et al., 2016; Easwar et al., 2023, using ERPs). At stimulus intensities well above the (unaided) hearing threshold, differences between conditions were smaller or even absent in the literature (Korczak et al., 2005; Vanheusden et al., 2020, using ERPs and neural tracking). Therefore, we believe that enhancements in neural tracking will decrease as stimulus intensity increases, in line with the first hypothesis. Lastly, we hypothesize that, similar to previous studies (e.g. Vanthornhout et al., 2018; Lesenfants et al., 2019; Van Hirtum et al., 2023), neural tracking of the speech envelope is significantly correlated with behaviorally measured speech intelligibility. Altogether, these investigations may provide an important step leading to a new objective EEG-based tool to evaluate hearing function and hearing aid benefit in hearing-impaired children.

## Material and Methods

### Participants

In total, ten children (6 female and 4 male) with bilateral mild to moderately severe sensorineural hearing loss participated in the present study. Eight children were included in experiment 1 (mean age: 7;02, range: 4;07-9;09 years), and eight children participated in experiment 2 (mean age: 8;04, range: 5;01-10;02 years). Six out of the ten children (S01-S06) included in this study participated in both experiments. Hearing thresholds were obtained by the children’s audiologist prior to the study. All children were experienced hearing aid users who had been using their hearing aids regularly for more than four years and had normal outer and middle ear function. Additionally, all children were native Dutch speakers, followed mainstream education, and had normal or corrected to normal vision. Relevant demographic features are given in Table 1. The study was approved by the Medical Ethics Committee UZ Leuven / Research (KU Leuven) (reference no. S57102) and all parents provided written informed consent before the experiment. After the experiment all children received a gift voucher to thank them for participating.

**Table 1.** Relevant participant details

| Subject | Age<br>experiment 1 | Age<br>experiment 2 | Right ear<br>PTA (dB) | Left ear<br>PTA (dB) | Degree of<br>hearing loss |
| --- | --- | --- | --- | --- | --- |
| S01 | 8;09 | 9;03 | 49 | 48 | moderate |
| S02 | 9;09 | 10;02 | 35 | 43 | moderate |
| S03 | 9;09 | 10;02 | 34 | 35 | mild |
| S04 | 6;06 | 7;02 | 60 | 62 | moderately severe |
| S05 | 6;10 | 7;04 | 44 | 43 | moderate |
| S06 | 4;07 | 5;01 | 43 | 45 | moderate |
| S07 | 5;07 | n/a | 64 | 54 | moderately severe |
| S08 | 5;06 | n/a | 42 | 46 | moderate |
| S09 | n/a | 8;01 | 75 | 68 | severe |
| S10 | n/a | 9;09 | 63 | 53 | moderate |

*Note.* Age is expressed in years;months.

### Behavioral experiment

Speech intelligibility was assessed using the Leuven Intelligibility Peuter Test (Lilliput; van Wieringen and Wouters, 2022). The lilliput contains 20 lists of 11 consonant-vowel-consonant words (e.g. “bus”) uttered by a female speaker. The number of phonemes correctly repeated by the child is counted to calculate the final outcome measure (i.e. the phoneme score). All lists were presented through a GENELEC (8020A) loudspeaker positioned at head-height of the seated child using the software platform APEX (Francart et al., 2008) on a Samsung Galaxy Tab A tablet. Children were instructed to look at the loudspeaker and to repeat each word or sound as accurately as possible. Each child started with a training list at 70 dB A with their personal hearing aids set according to the child’s usual prescribed settings. Next, multiple lists were presented at different intensities ranging between 30 and 80 dB A. First, a minimum of three lists representing soft, normal and loud speech were presented with hearing aids, followed by at least three lists without hearing aids. If necessary, additional lists were presented at a higher or lower intensity to ensure we obtained a score around 80% or higher (with 80 dB A as a maximum intensity), a score close to 50% and a score below the 50%-point. The behavioral test was conducted before the EEG measurement in both experiment 1 and experiment 2.

### EEG experiment

#### Speech material and procedure

**Experiment 1** In the first experiment, children listened to four different stories of “Little Polar Bear” during the EEG recordings. These stories are part of the children’s series by Hans de Beer, and were narrated by the same native Flemish speaker as the Lilliput. All stories were 10 to 12 minutes long and presented in two-minute fragments. In accordance with the behavioral experiment, speech was presented at a minimum of three intensities representing soft (40 dB A), normal (60 dB A) and loud (80 dB A) speech. Every two-minute fragment was presented randomly at a different intensity level, and each level was presented in an aided (i.e. with hearing aids) and unaided (i.e. without hearing aids) condition. In addition, the order in which the stories were presented was randomized. The aided condition was always presented first to optimize attention and motivation. In total, children listened to 6 minutes of speech (i.e. three two-minute fragments) at each level per condition (i.e. aided and unaided), resulting in at least 6 measurement conditions (*∼*36 minutes) for each child. For four subjects an additional 6 minutes were presented in the unaided condition at 50 dB A. For one subject, only 4 measurements could be carried out due to a limited attention span. An overview of the different blocks within an example test session is shown in Figure 1 (upper panel). While listening to the stories, children watched images from the “Little Polar Bear” books being projected on a sound-permeable screen in front of them to mimic a storytelling environment and to keep them engaged even when the story was not audible (e.g. at 40 dB A without hearing aids).

**Figure 1.**
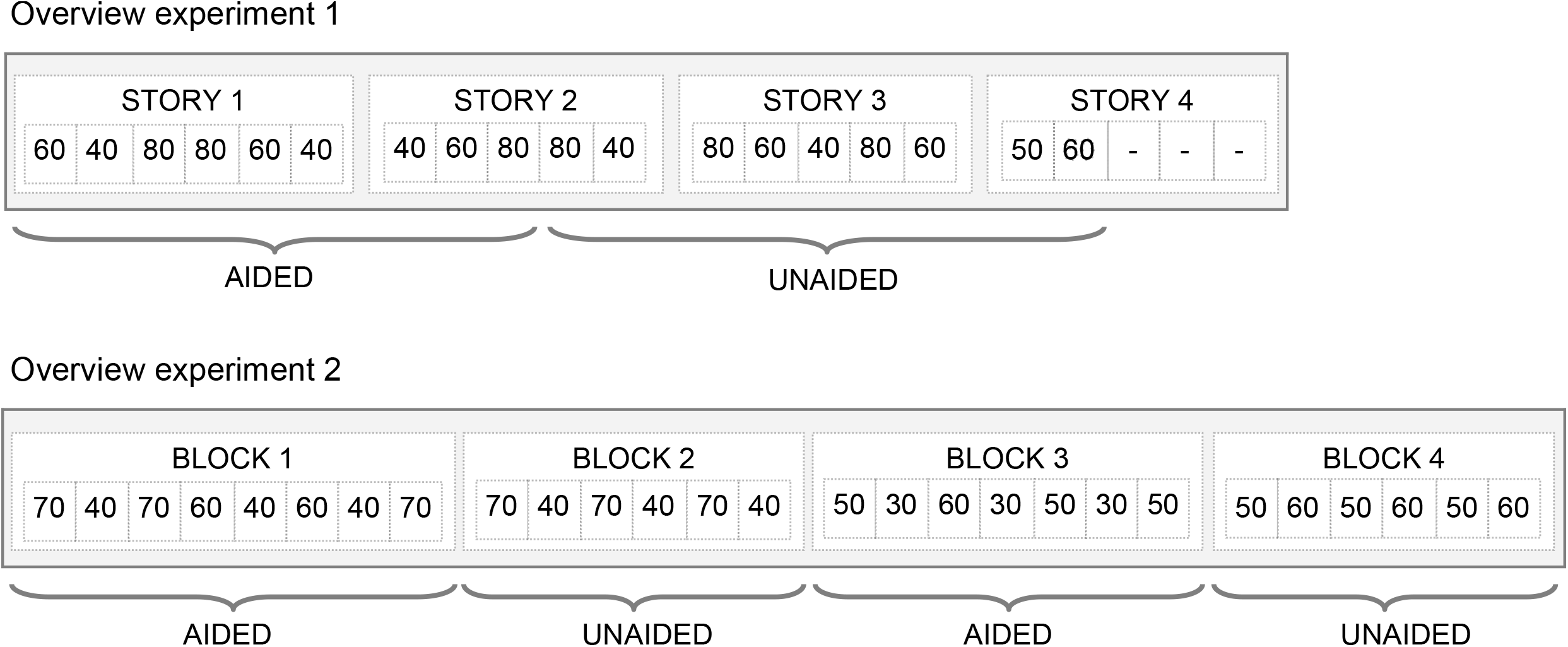
Overview of an example EEG test session for experiment 1 (upper panel) and 2 (lower panel). Each session started with the behavioral experiment where speech intelligibility was measured using the Lilliput. The numbers in each block indicate a 2-minute segment of the presented stimulus intensity in dB A. Every intensity was presented three times per condition (aided and unaided) to obtain 6 minutes of speech. Each child thus listened to 36 minutes of speech during experiment 1, and 54 minutes of speech during experiment 2. Because of the longer test duration of experiment 2, two (or three) stimulus intensities were grouped together in one block and presented first aided, and then unaided. This was done to ensure we would have enough data to compare aided and unaided conditions, in case a child would not be able to complete the full protocol.

**Experiment 2** Although the narrated stories used in the first experiment are naturally spoken stories, they do not capture some aspects of real-world listening. Therefore, in experiment 2, we used an animated movie which included more dynamic speech and naturalistic dialogues, without adding visual articulatory cues. The presented movie was “The Big Bad Fox and Other Tales” by Benjamin Renner and Patrick Imbert, narrated by different native Flemish speakers (male, female and children). The movie was 58 minutes long and presented in 4 blocks of two-minute segments. Based on the results of experiment 1, we decided to enlarge the range of stimulus intensities for the measurements of experiment 2. Stimulus intensities were fixed and ranged from loud to soft in steps of 10 dB: 70, 60, 50, 40 and 30 dB. Each level was presented in an aided and unaided condition, with the exception of 30 dB (only aided). The presentation order was semi-randomized. That is, each block contained two (or three) different intensities, combining an easier (e.g. 70 dB) and a more difficult (e.g. 40 dB) presentation level to keep children motivated and limit fatigue. Within one block, those intensities were presented three times, to obtain 6 minutes of speech at the same stimulus intensity. Additionally, hearing aids were always on or off within one block (i.e. hearing aid condition was fixed) to prevent taking the hearing aid on and off every 2 minutes. An overview of the different blocks within an example test session is shown in Figure 1 (lower panel). In total, children listened to 6 minutes of speech (i.e. three two-minute fragments) at each level per condition (five levels aided; four levels unaided), resulting in 9 measurement conditions (*∼*54 minutes) per child. For three children, only 8 measurement conditions are available due to restricted attention. There was a minimum of 5 and a maximum of 8 months between experiment 1 and experiment 2.

#### Data acquisition

All EEG recordings were conducted in a sound-attenuating, electrically shielded booth using a child-friendly and age-appropriate protocol. Children were seated in a comfortable chair, approximately 1 m in front of a sound-permeable screen and loudspeaker. The speech material was presented at a sample rate of 48000 Hz through a GENELEC (8020A) loudspeaker positioned at head-height of the seated child using the software platform APEX (Francart et al., 2008; experiment 1) or in-house stimulation software (experiment 2), and an RME Fireface UC soundcard (Haimhausen, Germany).

EEG data was recorded at a sampling rate of 8192 Hz, using a Biosemi ActiveTwo System with 64 Ag/AgCl electrodes based on the international 10-20 system. Electrode offsets were kept within the −30mV and 30mV range to ensure stable recording.

All children were instructed to attend to the presented stimulus (i.e. stories or movie), while minimising movements. To ensure compliance from the children and monitor alertness and movement, an experienced researcher sat alongside the child during the recording. Short recesses between each two-minute segment and each different story (or measurement block) were used to motivate the children, and if the child became too inattentive, fussy or tired, they were allowed to rest before resuming the recording.

### Signal processing

#### EEG pre-processing

All analyses were conducted offline using Matlab R2016b (The MathWorks Inc, 2016). First, the EEG data was downsampled from 8192 Hz to 256 Hz, to decrease computation time. Common EEG artifacts, such as eye blinks and muscle artifacts, were removed using a multi-channel Wiener filter (Somers et al., 2018). Next, bad EEG channels were interpolated using the five nearest, neighbouring channels channels using Fieldtrip (Oostenveld et al., 2011), before all channels were re-referenced to a 64-channel common-average. Finally, the EEG data were band-pass filtered between 0.5 and 4 Hz (delta band) using a Chebyshev filter (with 80 dB attenuation and 10% outside the passband), before being downsampled to 128 Hz.

#### Envelope reconstruction

In order to assess neural envelope tracking, we used an envelope reconstruction approach as described in more detail in by Vanthornhout et al. (2018) and Verschueren et al. (2020). In short, this involved extracting the speech envelope using a gammatone filter bank followed by a power law. The resulting acoustic speech envelope was then downsampled from 48000 to 256 Hz to reduce the computational load. Next, the speech envelope was band-pass filtered between 0.5 and 4 Hz (delta band) similarly to the EEG data. Lastly, the speech envelope was further downsampled to 128 Hz. To obtain the reconstructed envelope for each 6-minute condition (i.e. every stimulus intensity, with and without hearing aid), a condition-specific linear decoder was applied per subject. These decoders were calculated using ridge regression, as implemented in the mTRF toolbox (Lalor et al., 2006). Essentially, each decoder acts as a spatiotemporal filter that combines the 64 EEG channels and their time-shifted versions from 0 to 250 ms (the integration window) to generate a reconstruction of the envelope. Following normalization, envelope decoders were trained through a leave-one-out 6-fold cross-validation scheme. That is, for each condition, which included 6 minutes of EEG data, 5 minutes were utilized to train the decoder. The trained decoder was then applied on the remaining minute of EEG data to generate a 1-minute envelope reconstruction. This procedure was repeated 6 times to produce envelope reconstructions for all folds. Then, all envelope reconstructions within each condition were concatenated for further analysis. To calculate an envelope tracking value (i.e. the reconstruction accuracy) per condition, the original speech envelope was compared to the reconstructed envelope using a bootstrapped Spearman correlation. To determine the significance of this correlation value, a null distribution was constructed by correlating 1000 random permutations of the real and reconstructed envelopes with each other. The 95th percentile (one-tailed) of this distribution was taken to obtain a 95% confidence interval.

#### Temporal response function estimation

The envelope reconstruction approach described above integrates neural activity over multiple EEG channels and their time-shifted versions to reconstruct the envelope. However, this method fails to provide information concerning the spatial distribution of the response across the scalp. To overcome this limitation, and to evaluate the underlying neural processing separately for the aided and unaided conditions (which may not be similar), we conducted a linear forward modelling approach. In this approach, we predict the EEG given the acoustic speech envelope, resulting in a temporal response function (TRF) for each channel. A TRF represents an impulse response function that indicates how the brain reacts to the speech envelope. Estimating TRFs offers the advantage of providing insights into neural response latency, amplitude, and topography across the scalp. The initial signal processing steps were identical to those of the envelope reconstruction approach, with the only difference being that band-pass filtering is performed within a broader frequency band of 0.5-25 Hz. TRFs were computed using a 6-fold cross-validation for every condition, using 5 minutes for training and 1 minute for validation. The boosting algorithm (David et al., 2007) was used to compute TRFs for every channel, as described by the implementation in the Python Eelbrain toolbox (Brodbeck, 2020). In summary, boosting is an iterative algorithm, starting from a TRF model consisting of zeros, which is then improved iteratively by adding weights at a latency that results in the most significant enhancement in response prediction. The iterations stop when the model reaches a point where no further significant improvements can be made for a given step size by which the weights are modified. The TRFs were computed over an integration window between −200 and 500 ms using an adaptively decreasing step size. For further analysis and visualization, the estimated TRFs were convolved with a Gaussian kernel of 9 samples long (SD=2) to smooth over time lags. Finally, the smoothed TRFs were averaged across all folds.

### Statistical analysis

Statistical analysis were performed using R (version 4.0.3; R Core Team, 2020). All tests were performed with a significance level of *α* = 0.05 unless otherwise stated.

For the behavioral experiment, the speech reception threshold (SRT, i.e. the level at which 50% of the phonemes are intelligible), and the slope were determined by fitting psychometric functions to the phoneme scores for each child individually, using the following formula:

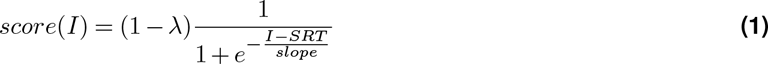

where “I” represents the stimulus intensity and *λ* the lapse-rate. For the behavioral experiment, *λ* was fixed to 0. We assessed the relation between neural envelope tracking, stimulus intensity and hearing aid condition by constructing a linear mixed effect model (LME) using maximum likelihood criteria with the nlme package (Pinheiro et al., 2022). Here, neural tracking is defined as the Spearman correlation between the real and the reconstructed envelope, and fixed and interaction effects of stimulus intensity and hearing aid condition (aided/unaided) are included. Additionally, a random intercept per participant was included in the model to account for dependencies between measures from the same child and to allow neural tracking to be higher or lower per participant. The residual plots of the final models were analysed to confirm the assumption of normality. All significant effects are discussed in the results section by reporting the *β* estimates, the corresponding standard error (SE), degrees of freedom (df), and test statistics (*t*-value and *p*-value). In addition, peak amplitude and latency of the TRF were determined for each individual child, by selecting the maximum TRF amplitude per intensity and condition between 50 and 130 ms. The relation between stimulus intensity and amplitude or latency was investigated by means of a Spearman correlation analysis per hearing aid condition. Additionally, the objective counterpart of the SRT, hereafter called the objective SRT, was estimated on the data across all subjects using the same formula to estimate behavioral SRTs, similar to the procedure described in detail by Vanthornhout et al. (2018). The score was calculated as the Spearman correlation between the actual and the decoded speech envelope.

To gain insight in the relation between neural envelope tracking and speech intelligibility, we performed a Spearman correlation analysis to assess the linear relation between envelope reconstruction scores and behaviorally measured speech intelligibility. If no behavioral score was available for a certain condition, we used the corresponding percentage-correct score from the individually fitted performance intensity functions for the correlation analysis.

## Results

### Experiment 1

#### Behavioral speech intelligibility

The individual unaided (diamonds) and aided (circles) SRTs are illustrated in Figure 2. The group-averaged unaided SRT was 54.77 dB A (SD=9.46) and ranged from 45.0 to 65.1 dB A. The group-averaged aided SRT was 37 dB A (SD=6.41) and ranged from 28.0 to 46.9 dB A.

**Figure 2.**
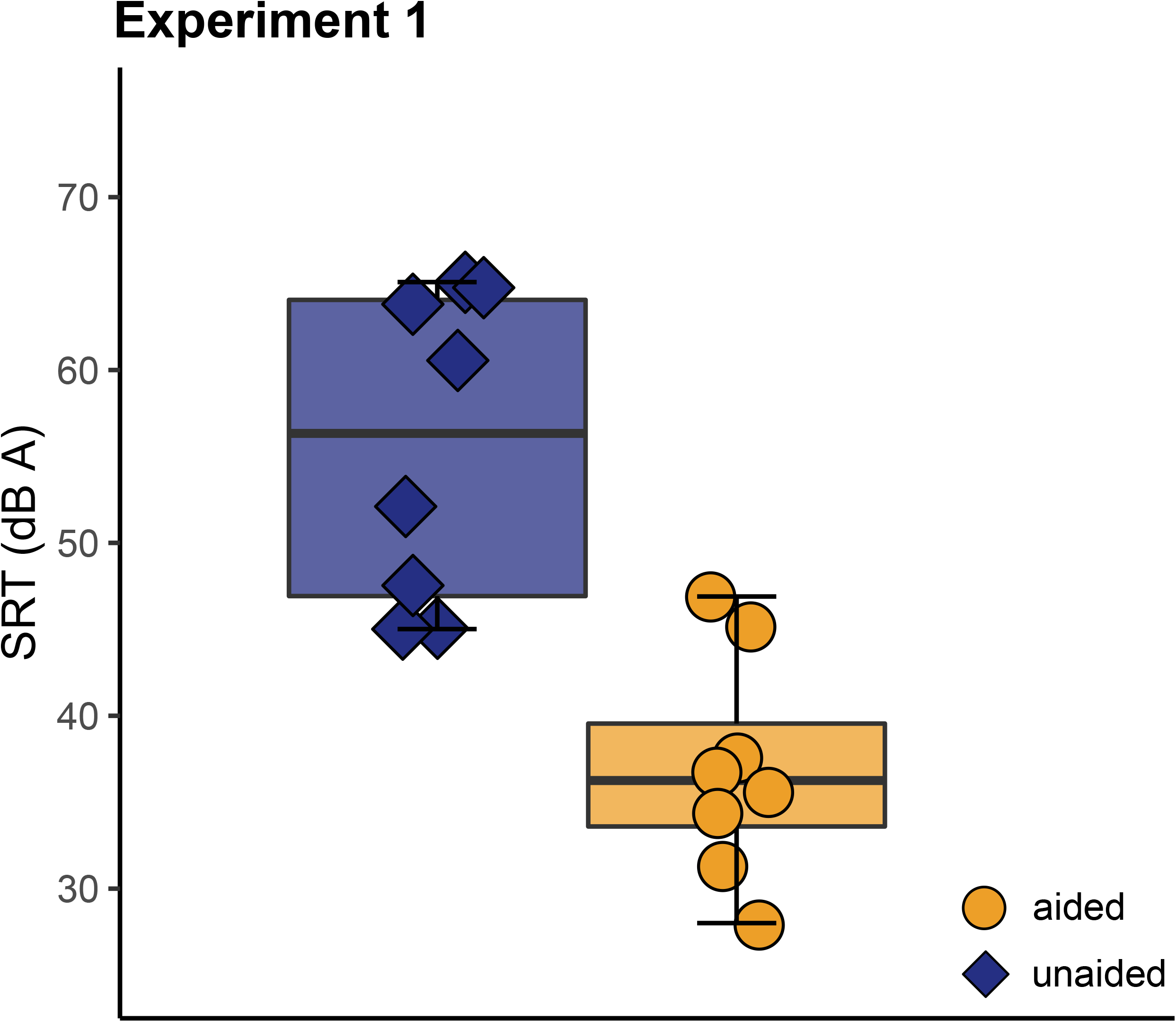
Speech reception thresholds of experiment 1. The boxplots show the distribution of the unaided (left, blue) and aided (right, orange) SRTs. Individual data points in each boxplot represent individual children, to show variation.

The individual intelligibility scores for each condition and the fitted psychometric functions from which the SRT was derived can be found in the Supplementary Information (see Figure S1).

#### Neural envelope tracking when listening to stories

First, we investigated the effect of stimulus intensity and hearing aid condition on neural envelope tracking based on the data obtained in experiment 1. Figure 3A shows the envelope reconstruction accuracies per child and condition at the different stimulus intensities. A linear mixed effect model was constructed with envelope reconstruction accuracy as the dependent variable, and factors hearing aid condition (unaided/aided) and stimulus intensity as fixed effects with random intercepts for each child. The model results (see Table S1) showed that the use of a hearing aid significantly improved envelope reconstruction accuracies, as expected (*b*=-0.3, SE=0.057, *p<*0.001). No significant effect of stimulus intensity (*b*=0.0002, SE=0.001, *p*=0.753) was found. However, the interaction term between hearing aid condition and stimulus intensity was significant (*b*=0.004, SE=0.001, *p<*0.001) as visualised in Figure 3B. This indicates that neural tracking increased significantly with increasing stimulus intensity, but only for the unaided condition (*b*=0.004, SE=0.001, *p<*0.001). No intensity-related difference was found in aided neural tracking (*b<*0.001, SE=0.001, *p*=0.773).

**Figure 3.**
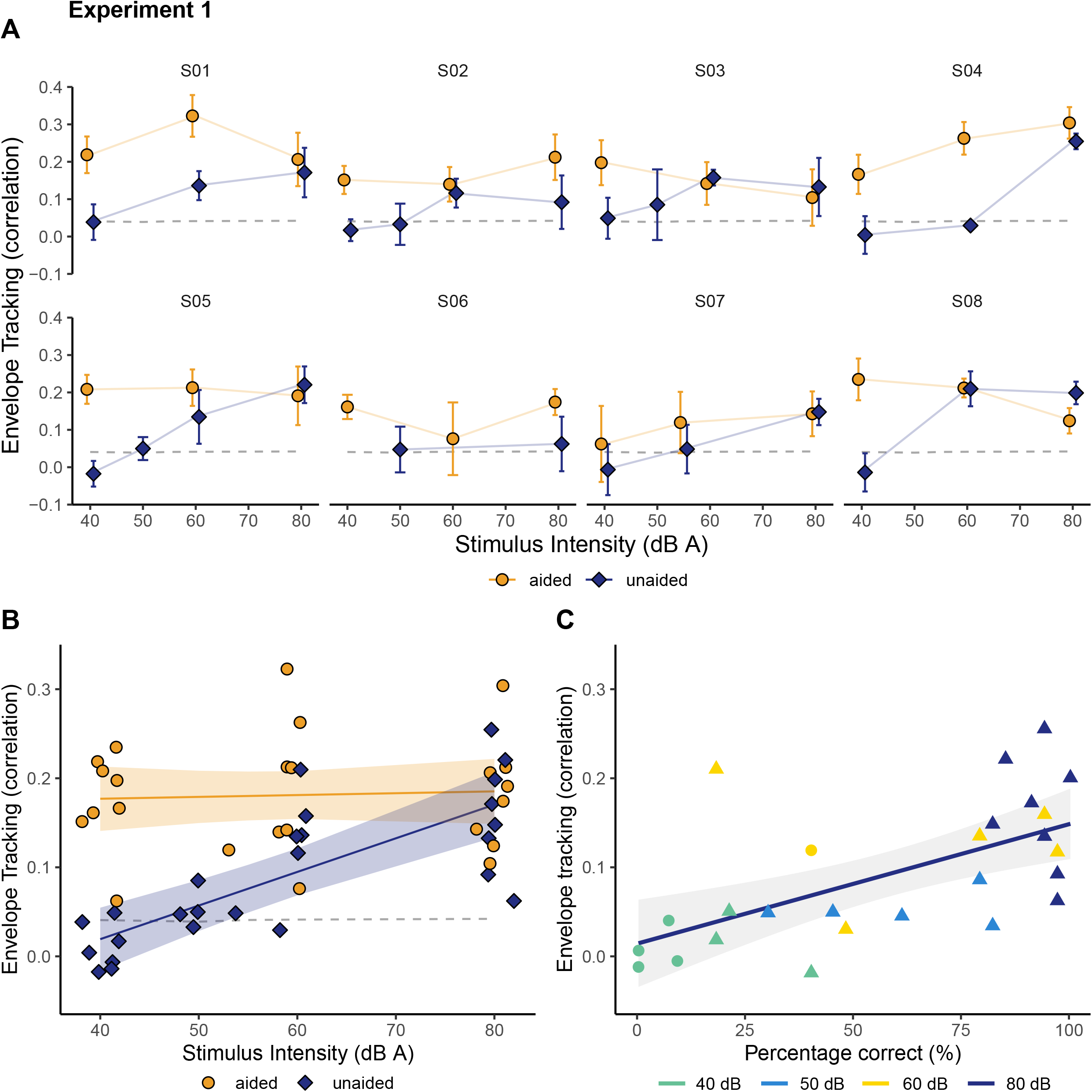
Envelope reconstruction accuracy as a function of stimulus intensity or speech intelligibility (Experiment 1). **A**. Neural envelope tracking for each individual child as a function of stimulus intensity. The error bars represent the 95% confidence level interval across all separate 1-minute folds. The dashed line (light grey) shows the significance level (i.e. the 95th percentile of the null distribution) of the envelope reconstruction scores. **B**. Neural envelope tracking as a function of stimulus intensity. The solid lines represents a linear model that was fitted on the data. The shaded area indicates the 95% confidence interval for the model predictions, and the significance level of the envelope reconstruction accuracy is indicated as the dashed line (light grey). **C**. Envelope tracking as a function of behavioral speech intelligibility in the unaided condition. Individual speech intelligibility scores are marked by colored triangles. If no datapoint was available, the corresponding percentage-correct score was determined from the individually fitted performance intensity functions (circles). The solid line shows the linear model fit. The shaded area represents the 95% confidence interval for the predictions of the linear model.

To further investigate whether neural tracking reflects speech intelligibility, rather than just changes in stimulus intensity, Spearman correlations between reconstruction accuracies and speech intelligibility scores were calculated. Figure 3C shows that neural tracking for the unaided condition increased with increasing speech intelligibility (*r* =0.59; *p*=0.002), suggesting that the better a child can understand speech, the higher the reconstruction accuracy for that child. We did not find a significant correlation between aided neural tracking and speech intelligibility (*r* =0.09; *p*=0.661). This result is not surprising considering the minimal fluctuations in behavioral scores, which indicate near-ceiling performance. That is, the majority of phoneme scores ranged between 80% and 100%).

Next, the spatiotemporal properties of the responses were investigated through forward modeling, as shown in Figure 4. We selected 12 frontocentral channels (presented by the black dots in Figure 4B) based on (Van Hirtum et al., 2023), and averaged them per subject, resulting in one TRF per intensity and condition for each subject. Figure 4A shows 2 prominent peaks appearing between 50-100 ms (Peak 1) and 150-200 ms (Peak 2). The topography of these peaks is shown in Figure 4B. The response activity at the peak latencies is predominantly fronto-centrally located and appears stronger for aided conditions compared to unaided. In the aided condition, when the stimulus intensity decreases, there appears to be an increase in the latency of peak 1. In the unaided condition, peak 1 completely disappears when the stimulus intensity is 40 dB, and speech understanding is limited (i.e. *<*20%). On the other hand, peak 2 can be discerned at later latencies, but only in response to the highest stimulus intensity (i.e. 80 dB A). Therefore, more detailed analysis was only carried out for Peak 1. Peak amplitude and latency were individually determined by selecting the maximum TRF amplitude per intensity and condition between 50 and 130 ms. Then, we investigated the amplitude and latency of these individual peaks as a function of stimulus intensity per condition (see Table S2-S5) As expected, in the unaided condition the amplitude of Peak 1 increased with increasing stimulus intensity (*b*=0.0003, SE=0.0001, *p*=0.005), whereas the latency of peak 1 did not show a significant relation with stimulus intensity (*b*=-0.204, SE=0.308, *p*=0.519). In contrast, for the aided condition the latency of peak 1 decreased with increasing stimulus intensity (*b*=-0.565, SE=0.189, *p*=0.009), whereas the amplitude of peak 1 did not change significantly as a function of stimulus intensity (*b<*0.001, SE*<*0.001, *p*=0.632), similar to the envelope reconstruction analysis. Scatterplots and regression lines showing the associations between the peak characteristics and stimulus intensity can be found in the Supplementary Information (see Figure S2).

**Figure 4.**
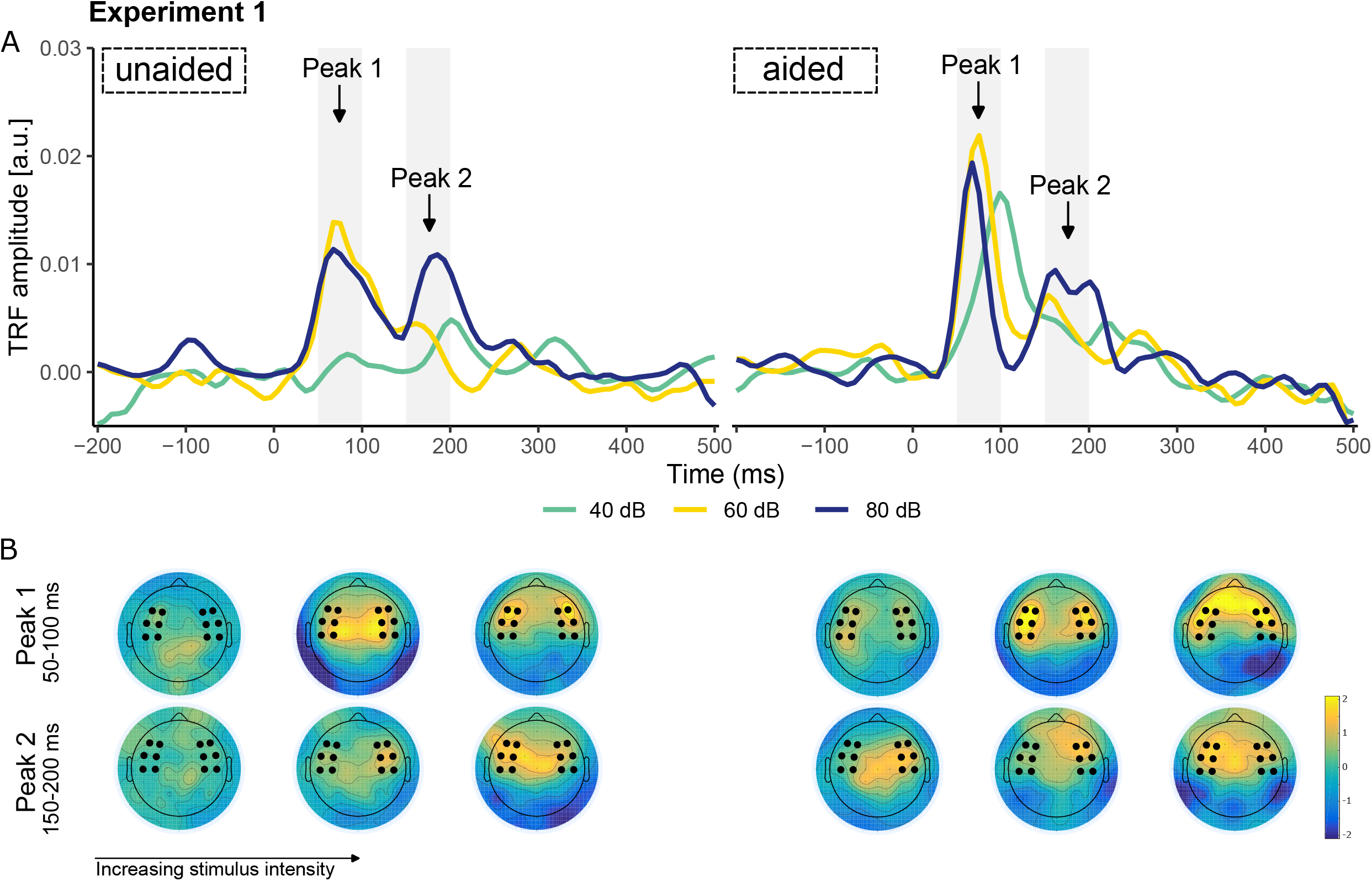
The effect of stimulus intensity and hearing aid condition on spatio-temporal properties of envelope tracking in experiment 1. **A**. Mean temporal response functions (TRF) across participants per stimulus intensity. **B**. Topographies showing mean TRF activity at the latencies of peak 1 (50-100 ms) and peak 2 (150-200 ms). The fronto-central electrode selection over which the TRFs in panel A were averaged is indicated by the black dots.

### Experiment 2

#### Behavioral speech understanding

The individual unaided (diamonds) and aided (circles) SRTs are illustrated in Figure 5. The group-averaged unaided SRT was 51.81 dB A (SD = 11.16) and ranged from 39.5 to 73.0 dB A. The group-averaged aided SRT was 34.67 dB A (SD = 8.83) and ranged from 26.7 to 49.9 dB A. As expected, given that most participants (N=6) participated in both experiments, these results are in line with the results found in the first experiment.

**Figure 5.**
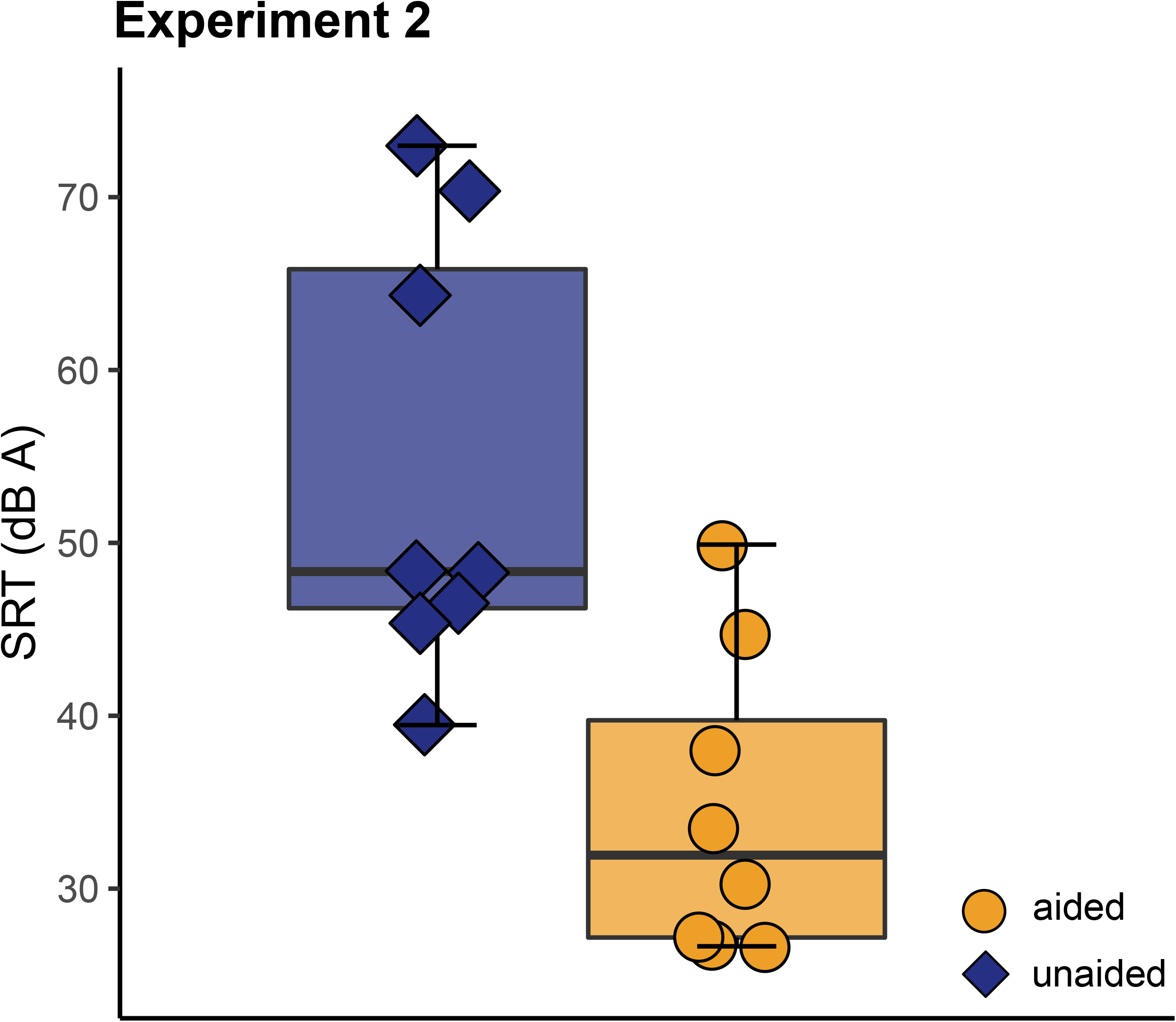
Speech reception thresholds of experiment 2. The boxplots show the distribution of the unaided (left, blue) and aided (right, orange) SRTs. Individual data points in each boxplot represent individual children, to show variation.

The individual intelligibility scores for each condition and the fitted psychometric functions from which the SRT was derived can be found in the Supplementary Information (see Figure S3).

#### Neural envelope tracking when watching an animated movie

Given that aided speech intelligibility at 40 dB A still exceeded 50% in experiment 1, the range of stimulus intensities presented in experiment 2 was enlarged, with the lowest intensity (in the aided condition) fixed at 30 dB A. This would allow us to investigate potential changes in aided neural tracking, that would reflect a decrease in speech intelligibility over and beyond decreasing stimulus intensity. Figure 6A shows the envelope reconstruction accuracies per child and condition as a function of stimulus intensity. A linear mixed effect model was constructed, identical to the analysis carried out for experiment 1. As illustrated in Figure 6B, we found an increase of neural tracking with increasing stimulus intensity (*b* = 0.002, SE *<* <0.001, *p <* 0.001, see Table S6) and a clear benefit of hearing aid on the envelope reconstruction accuracies (*b* = −0.296, SE = 0.058, *p <* 0.001). Additionally, the significant interaction between stimulus intensity and hearing aid condition, as found in the first experiment, was also apparent in experiment 2 (*b* =0.004, SE = 0.001, *p <* 0.001), showing that hearing aid benefit increased as intensity decreases (*b* = −0.006, SE = 0.001, *p <* 0.001). Moreover, neural tracking increased significantly with increasing stimulus intensity for both the aided and unaided condition. However, when excluding the 30 dB A condition from the analysis for the aided condition, no significant effect of stimulus intensity on envelope reconstruction accuracy could be found (*b* = 0.001, SE = 0.001, *p* = 0.237).

**Figure 6.**
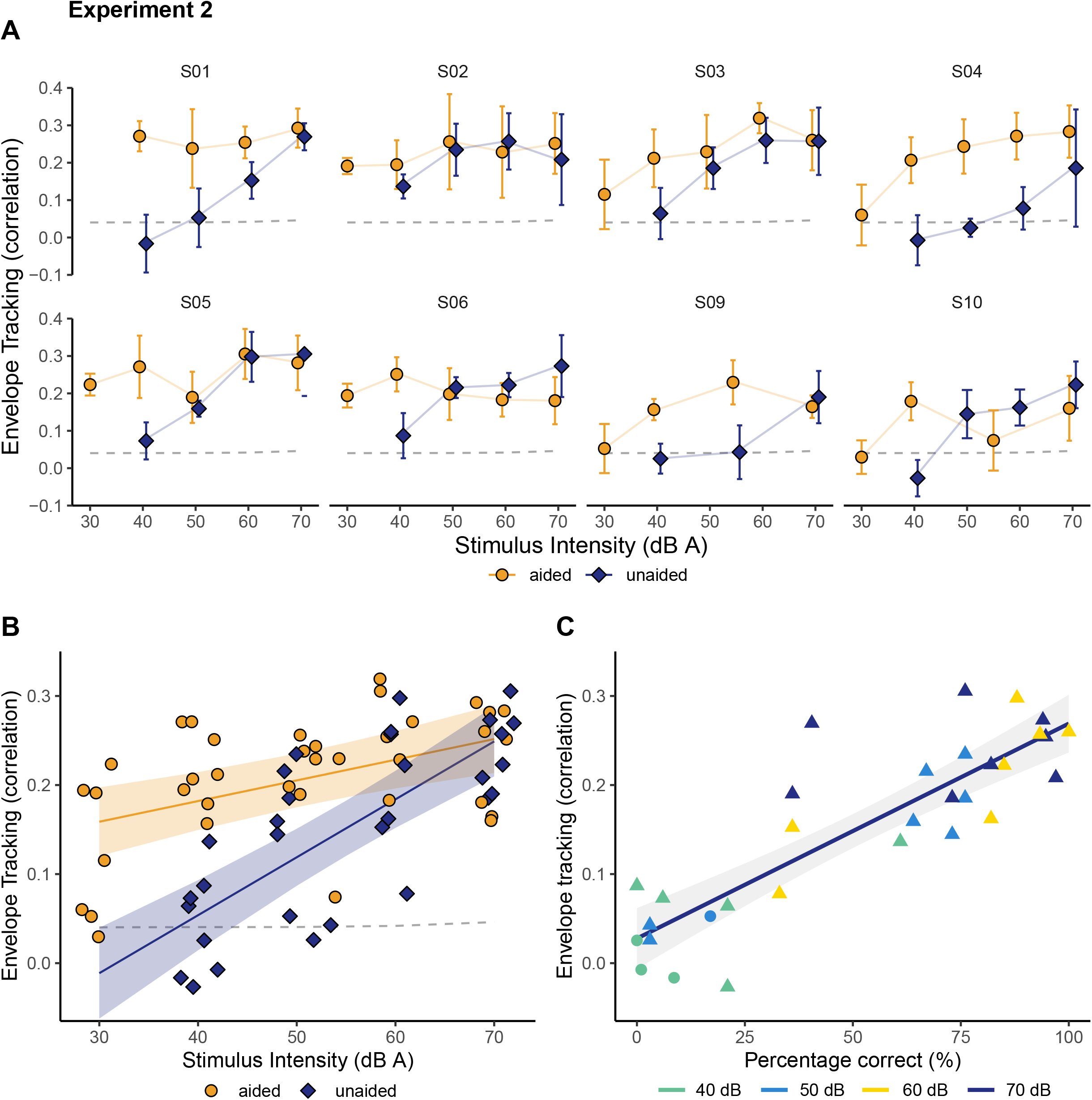
Envelope reconstruction accuracy as a function of stimulus intensity or speech intelligibility (Experiment 2). **A**. Neural envelope tracking for each individual child as a function of stimulus intensity. The error bar represents the 95% confidence level interval across all separate 1-minute folds. The dashed line (light grey) shows the significance level (i.e. the 95th percentile of the null distribution) of the envelope reconstruction scores. **B**. Neural envelope tracking for aided (orange, circles) and unaided (blue, diamonds) conditions as a function of stimulus intensity. The solid lines represents a linear model that was fitted on the data. The shaded area indicates the 95% confidence interval for the model predictions, and the significance level of the envelope reconstruction accuracy is indicated as the dashed line (light grey). **C**. Envelope tracking as a function of speech intelligibility in the unaided condition. Individual speech intelligibility scores are marked by colored triangles. If no datapoint was available, the corresponding percentage-correct score was determined from the individually fitted performance intensity functions (circles). The color gradient was used for illustrative purposes to mark stimulus intensity. The solid line shows the linear model fit. The shaded area represents the 95% confidence interval for the predictions of the linear model.

The relation between neural tracking and behavioral speech intelligibility was again quantified by calculating Spearman correlations between reconstruction accuracies and speech intelligibility scores for each condition. The results obtained further support the findings from experiment 1. As depicted in Figure 6C, neural tracking in the unaided condition increased as speech intelligibility scores improved (*r* =0.83; *p<*0.001). Furthermore, we found a significant positive correlation between aided neural tracking and speech intelligibility (*r* =0.63; *p<*0.001). Again, this relation disappears when excluding the 30 dB A level (*r* =0.33; *p*=0.078), suggesting that neural tracking correlates with speech intelligibility rather than stimulus intensity.

Finally, we calculated TRFs for every 6-min condition. The final averaged TRFs per condition are shown in Figure 7A. A first peak (Peak 1) can be discerned between 50 and 100 ms for both the aided and unaided condition. Furthermore, a distinct second peak emerges at later latencies, specifically between 150 and 200 ms, although it is more prominently visible in the aided condition. The topography of these peaks is shown in Figure 7B and has a similar pattern as the results found in the first experiment. That is, a predominantly fronto-centrally located response activity. At the later peak, the frontocentral activity is slightly reduced and appears more right dominant. The influence of stimulus intensity and hearing aid condition was investigated in more detail by determining the peak latency and amplitude of the first peak. As shown in Figure 7A, and confirmed by statistical analysis (see Table S7-S10), with increasing stimulus intensity, the amplitude of peak 1 in the unaided condition increases (*b*=0.0006, SE=0.0001, *p<*0.001), whereas the latency of peak 1 did not show a similar effect (*b*=0.179, SE=0.329, *p*=0.592). Consistent with the results of the first experiment, we did not find a significant correlation between the amplitude of peak 1 and stimulus intensity in the aided condition (*b*=0.0002, SE=0.0001, *p*=0.125). However, the latency of peak 1 decreased significantly with increasing stimulus intensity (*b*=-0.833, SE=0.129, *p<*0.001). Scatterplots and regression lines showing the association between the peak characteristics and stimulus intensity can be found in the Supplementary Information (see Figure S4).

**Figure 7.**
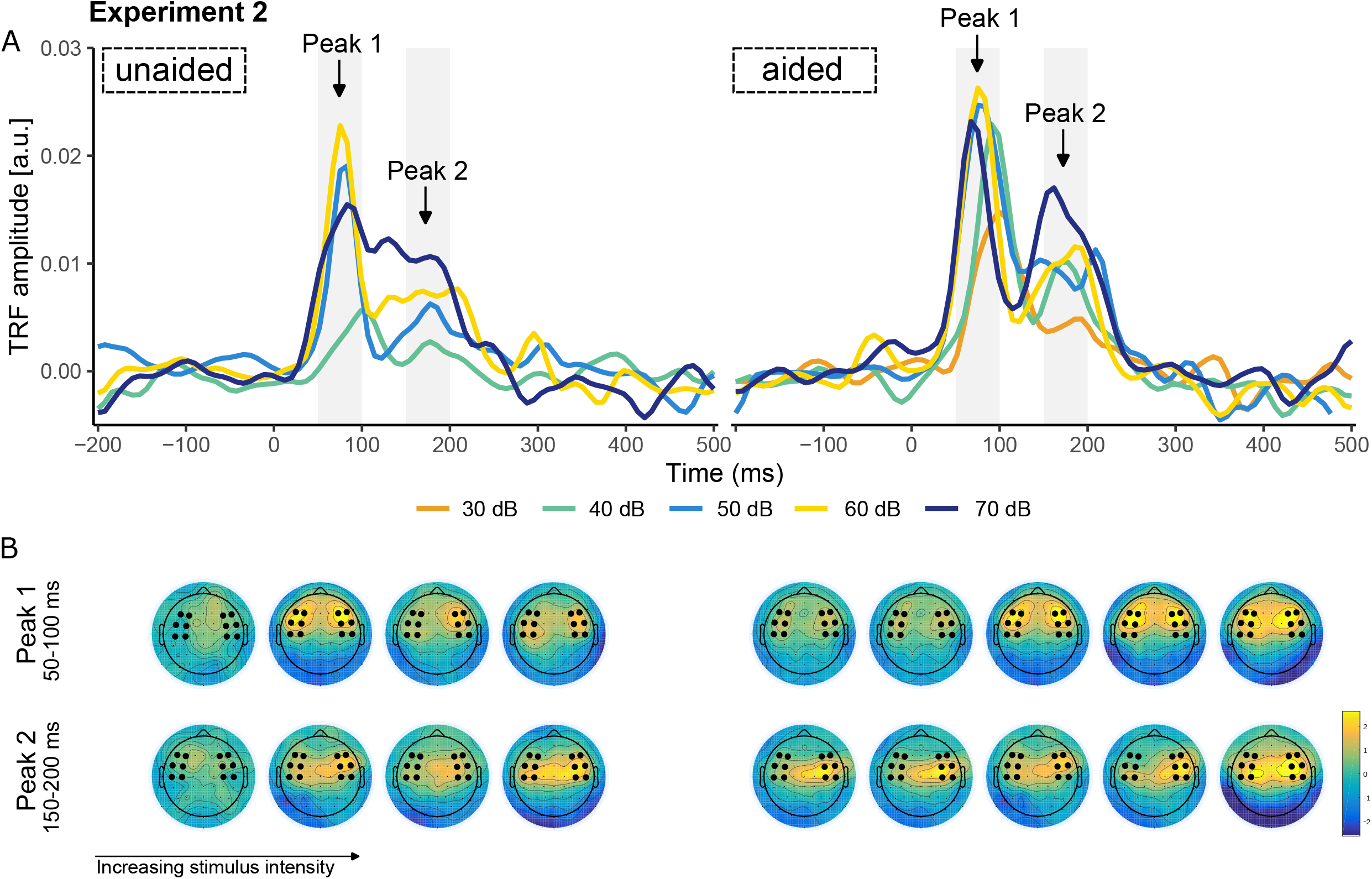
The effect of stimulus intensity and hearing aid condition on spatio-temporal properties of envelope tracking in experiment 2. **A**. Mean temporal response functions (TRF) across participants per stimulus intensity for the unaided (left) and aided (right) condition. **B**. Topographies showing mean TRF activity at the latencies of peak 1 (50-100 ms) and peak 2 (150-200 ms). The fronto-central electrode selection over which the TRFs in panel A were averaged is indicated by the black dots.

### The effect of stimulus choice

In the present study, two experiments were conducted. In the first experiment, neural tracking was assessed by presenting narrated stories accompanied by story illustrations, whereas in the second experiment, an animated movie was used as the stimulus. Although the outcomes of both experiments were overall identical, it is interesting to explore the potential influence of stimulus choice on neural envelope tracking. The data of the two experiments were combined for the children that participated in both experiments and a linear mixed effect model was computed with neural tracking (i.e. envelope reconstruction accuracy) as the dependent variable, and fixed and interaction effects of stimulus intensity, hearing aid condition and stimulus. The model results (see Table S11) showed an effect of stimulus intensity (*b* = 0.002, SE = 0.001, *p <* 0.001), hearing aid condition (*b* = −0.274, SE = 0.049, *p <* 0.001) and a significant interaction between stimulus intensity and hearing aid condition (*b* = 0.004, SE = 0.001, *p <* 0.001) as expected. In addition, the interaction term between stimulus intensity and stimulus was significant (*b* = −0.002, SE = 0.001, *p* = 0.007), suggesting that reconstruction accuracies were higher when presenting a movie compared to a story. Additionally, this enhancement becomes more apparent as stimulus intensity increases. On the other hand, the interaction term between stimulus and hearing aid condition was not significant (*b* = −0.041, SE = 0.025, *p* = 0.101), indicating that both narrated stories or animated movies could be used to evaluate hearing aid benefit. Next, we estimated the objective counterpart of the SRT from the neural tracking data in order to explore whether variations in reconstruction accuracies would correspond to differences in the objective SRT. As aided speech intelligibility generally exceeds 50% at the lowest presented intensity, we focused our analyses solely on the unaided condition. Since aided speech intelligibility is on average higher than 50% on the lowest intensity presented, the analyses were only carried out for the unaided condition. A sigmoid was fitted on the unaided reconstruction accuracies across all subjects for each stimulus separately. After fitting the function, we derived its midpoint, which we will refer to as the objective SRT. As illustrated in Figure 8, the objective SRT was 55.58 dB (A) for the story (Experiment 1) and 46.29 dB (A) for the movie (Experiment 2), suggesting that the movie might be easier to understand compared to the narrated stories.

**Figure 8.**
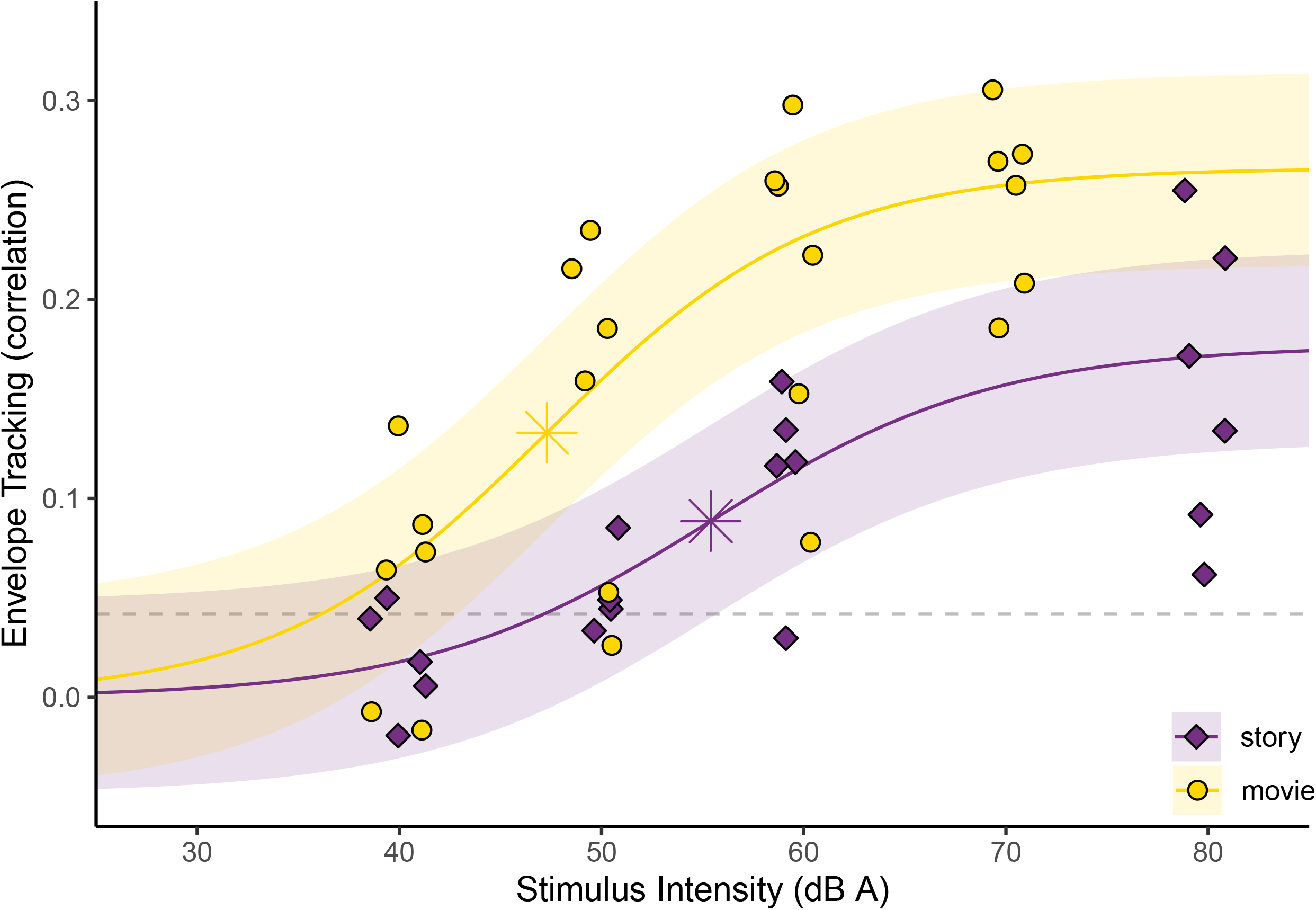
Comparison of neural envelope tracking using a story (purple, diamonds) or a movie (yellow, circles) as a stimulus for the unaided condition. The solid lines represents a sigmoid function that was fitted on the data. The star represents the estimated objective SRT for each stimulus. The shaded area indicates the root mean square error of the fit, and the significance level of the envelope reconstruction accuracy is indicated as the dashed line (light grey).

## Discussion

We investigated whether neural envelope tracking could serve as a potential indicator of hearing function (i.e. speech intelligibility) and hearing aid benefit in children with permanent hearing loss, by conducting two separate experiments. In the first experiment, we recorded EEG in 8 hearing-impaired children as they listened to age-appropriate stories in quiet, using multiple audible and inaudible stimulus levels. In the second experiment, we followed a similar protocol, but used an animated movie as the stimulus to better reflect real-life speech. The findings of both experiments were consistent and aligned with our hypotheses. Firstly, we observed that neural envelope tracking increased with increasing stimulus intensity in the unaided condition. In contrast, neural tracking remained stable in the aided condition, as long as speech intelligibility was maintained. Secondly, the use of personal hearing aids significantly improved neural envelope tracking. Furthermore, the benefit of hearing aids increased as stimulus intensity decreased and speech became more challenging to understand. Finally, we found a significant correlation between neural envelope tracking and behavioral measures of speech intelligibility.

### Relation between neural envelope tracking, stimulus intensity and speech intelligibility

We hypothesized that neural envelope tracking in the delta band would increase as a function of stimulus intensity, given that the changes in stimulus intensity cause changes in speech intelligibility. We demonstrated that this is indeed the case in the unaided condition, consistent with previous EEG work using mostly speech-in-noise paradigms (Vanthornhout et al., 2018; Decruy et al., 2019; Lesenfants et al., 2019; Iotzov and Parra, 2019; Etard and Reichenbach, 2019; Verschueren et al., 2020). As expected, in the aided condition, stimulus intensity did not significantly influence neural envelope tracking, at least not between 40 dB A and 80 dB A. Only by including a condition with a lower intensity of 30 dB A in the second experiment, we could observe an increase of neural tracking of the speech envelope with increasing stimulus intensity. These findings are corroborated by multiple studies suggesting that acoustically degrading the speech signal, by lowering the stimulus intensity or adding background noise, does not necessarily alter neural tracking, provided that speech remains intelligible (Ding and Simon, 2013; Verschueren et al., 2021; Van Hirtum et al., 2023). For example, Van Hirtum et al. (2023) showed an S-shaped (sigmoid) correlation across SNRs, indicating that as the SNR decreased, envelope reconstruction accuracy in the delta band remained stable until behaviorally measured speech intelligibility started to decrease.

The above-mentioned studies all featured normal-hearing adults or children. However, Decruy et al. (2020), for example, also found that neural tracking increased with increasing speech understanding in hearing-impaired adults, even after amplifying the presented speech stimuli using linear amplification. The authors utilized subject-specific SNRs for stimulus presentation, covering the psychometric function of each participant. This ensured that differences between conditions were solely attributed to differences in speech intelligibility. Additionally, Presacco et al. (2019) also noted an increase in neural envelope tracking correlated with SNR for hearing-impaired adults, but not for normal-hearing adults. This disparity could be attributed to the high intelligibility of all presented SNRs among the normal-hearing group, similar to our aided findings.

Like previous research, we also observed a significant correlation between speech intelligibility and neural envelope tracking in both normal-hearing (Meyer et al., 2017; Vanthornhout et al., 2018; Etard and Reichenbach, 2019; Verschueren et al., 2020, 2021; Van Hirtum et al., 2023), and hearing-impaired listeners (Verschueren et al., 2019; Decruy et al., 2020; Paul et al., 2020). These studies consistently report that increased neural envelope tracking indicates improved speech intelligibility. Furthermore, it has been found that neural envelope tracking is stronger when listening to intelligible speech compared to unintelligible signals, even after accounting for audibility (Ahissar et al., 2001; Doelling et al., 2014; Peelle et al., 2013; Gross et al., 2013; Di Liberto et al., 2018). Altogether, our findings suggests that neural tracking correlates with speech intelligibility rather than stimulus intensity or audibility, at least in the delta-band.

### Impact of personal hearing aids on neural envelope tracking

The use of amplification, by means of personal hearing aids, substantially enhances neural tracking. As hypothesized, the greatest benefit occurred for the lowest stimulus intensities, which would be inaudible without amplification. Only three neural tracking studies so far included hearing-impaired participants who were bilateral hearing aid users (Decruy et al., 2020; Mirkovic et al., 2019; Vanheusden et al., 2020). Notably, only the study of Vanheusden et al. (2020) had participants wear their personal hearing aids during testing, similar to our current study. Their results did not demonstrate any differences in neural tracking between aided and unaided conditions. However, speech was presented at a clearly audible and high intensity, resulting in good speech intelligibility even without hearing aids. This might explain why the hearing aid did not further enhance neural responses to the unaided speech envelope. This hypothesis aligns with observations of CAEPs, which indicate that the use of personal hearing aids has a significant impact on both the strength and timing of neural responses in infants (e.g. Chang et al., 2012) as well as adults (e.g. Van Dun et al., 2016). However, this effect is dependent on audibility of the stimuli, meaning that the increase in response amplitude is particularly evident at lower stimulus intensities (Korczak et al., 2005).

The results of the present study also show that the vast majority of hearing-impaired children consistently showed significant neural tracking when they wore their prescribed personal hearing aids. In the unaided condition, significant responses at lower intensities were limited. These observations align with the behavioral measures obtained. Specifically, aided percentage correct scores averaged over 50% at 30 dB and remained around 70% at approximately 40 dB. In contrast, percentage correct scores for the unaided condition were only about 20% at 40 dB and approximately 60% around 50 dB. Thus, we hypothesize that a change in the detection status, indicating whether neural tracking is significant or not, could serve as a clear indication of improved speech intelligibility with the use of a hearing aid. Additionally, the relative differences in reconstruction accuracy between the different conditions (i.e., the use of a hearing aid or higher stimulus intensity) can be utilized as a (diagnostic) marker for speech intelligibility(Gillis et al., 2022b).

### Effect of stimulus intensity and hearing aids on temporal response functions

We found two positive TRF peaks between 50-100 ms and 150-200 ms in the fronto-central region. These findings are generally consistent with previous research in adults (e.g. Ding and Simon, 2012a, 2013; Vanthornhout et al., 2019; Verschueren et al., 2021, 2022), although those studies suggested slightly faster occurrence of both peaks, with the first peak around 50 ms and the second peak between 100-150 ms. However, a recent study in 5-year-old children revealed a single prominent positive TRF peak around 80 ms (Van Hirtum et al., 2023), which aligns with the delayed latency of the first peak observed in our study. Additionally, in our study, we observed that the presence of the second peak was mostly evident at higher stimulus intensities, particularly at the individual level. The different TRF peaks are believed to represent distinct neural mechanisms involved in processing the stimulus over time. That is, the earlier peaks may primarily reflect acoustic processing, while the later peaks may involve more top-down driven processes like attention and comprehension (e.g. Brodbeck and Simon, 2020). On the other hand, the presence of a single peak in the TRF of young children, as observed by Van Hirtum et al. (2023), suggests that children’s neural activity exhibits a less differentiated response to the stimulus, which is consistent with previous observations of CAEPs (Ceponien et al., 1998; Albrecht et al., 2000; Ponton et al., 2000). Therefore, the difference in both the temporal pattern and latency between adults and children likely reflect developmental changes in the neural mechanisms and cognitive abilities involved in processing speech.

To further explore the impact of stimulus intensity and hearing aids om the TRFs, we compared individual peak amplitudes and latencies across different stimulus intensities. Our findings in the unaided condition revealed a positive correlation between stimulus intensity and the amplitude of the first peak (hereafter P1), which is consistent with prior research conducted on adults (e.g. Ding and Simon, 2013; Verschueren et al., 2021; Muncke et al., 2022 and children (Van Hirtum et al., 2023). These studies consistently observed a gradual decrease in the amplitude of the early TRF peak (*∼*50 ms for adults, *∼*80 ms for children) as a function of SNR or intensity. However, contrary to the aforementioned studies, we did not find an increase in latency with decreasing intensity. In contrast, the results in the aided condition showed an increase in P1 latency as the stimulus intensity decreased, while the P1 amplitude remained unaffected by the presentation level. To the best of our knowledge, there has been only one study examining TRFs in hearing-impaired listeners. Gillis et al. (2022a) observed a general trend of increased latency as the listening conditions became more challenging. However, for adults with hearing loss, this latency increase was not evident, which could possibly account for the absence of a distinct latency effect in our unaided results. Another possibility to consider is that the inability to find effects could be due to limited statistical power. Specifically, in the unaided condition, individual peaks are less distinguishable at lower intensities (e.g. 40 dB), where speech is either barely audible or no longer audible for most children. This leads to less reliable peak detection and significant variation in peak latency among subjects. On the other hand, in the aided condition, more reliable data points are available, making the latency effect more pronounced.

A delay in neural responses has been suggested to indicate a decline in neural processing efficiency, both in the neural processing of continuous speech (Bidelman et al., 2019; Gillis et al., 2022a; Verschueren et al., 2022), and simple sounds (Korczak et al., 2005; Billings et al., 2015; Maamor and Billings, 2017; McClannahan et al., 2019; Van Dun et al., 2016). Moreover, increased latencies have been associated with heightened task demands, such as lower stimulus intensity or elevated background noise (Mirkovic et al., 2019; Verschueren et al., 2021). Therefore, as stimulus intensity decreases, the task demand increases, requiring more processing time, which decreases neural processing efficiency. In our study, we observed the same effect of stimulus intensity in hearing-impaired children. However, the aided results in this study only showed significant differences in TRF latency, and not in TRF amplitude. This suggests that decreasing stimulus intensity affects neural response amplitude only at intensities where speech audibility impacts listening effort and/or speech intelligibility (Verschueren et al., 2021; Van Hirtum et al., 2023), such as in the unaided condition. Furthermore, despite improvements in audibility provided by personal hearing aids, the brain might not process speech with the same level of effectiveness as normal-hearing individuals. Therefore, we believe that TRF latency can serve as an indicator for the speech processing efficiency, while its amplitude and detectability provide important information about audibility of speech.

### Clinical implications

This study provides the first evidence that neural tracking of the speech envelope is correlated to speech intelligibility in hearing-impaired children. As for the actual testing procedure itself, the present study design demonstrates the practical feasibility of using natural, continuous speech stimuli to measure neural envelope tracking in children with and without hearing aids. For instance, our findings illustrate that both narrated stories and movie stimuli can be used to measure neural tracking. Nevertheless, we found differences between both stimuli. Specifically, neural tracking was enhanced when using the movie (experiment 2) compared to the stories (experiment 1). On a group level, we determined the objective counterpart of the speech reception threshold (SRT) based on the unaided reconstruction accuracies, which were 55.6 dB A for the story and 46.3 dB A for the movie, suggesting that the movie might be somewhat easier to understand compared to the narrated stories. Furthermore, a stronger correlation between speech intelligibility and neural envelope tracking was found when using an animated movie in experiment 2.

A potential confound in the study is that the stimuli were not matched in terms of speaker characteristics. Specifically, the stories were narrated by a single speaker, whereas the movie was narrated by different speakers. Additionally, the acoustics of the stimuli varied in terms of prosody and speech rate. Narrated stories may lack certain natural speech variations present in everyday communication, such as pitch, and often have a slower pace compared to natural conversations. Previous research has shown that these properties can influence neural tracking (Verschueren et al., 2022). Lastly, including an animated movie in this study was motivated by the fact that movies are more engaging for young children, especially when speech is inaudible. Considering that enhancements in neural tracking have been associated with the amount of contextual information in the stimulus (Verschueren et al., 2020), as well as attention (Lesenfants et al., 2019; Vanthornhout et al., 2019), it is possible that the differences observed between the two experiments can, at least partly, be attributed to these factors. Through observation, we indeed found that children were generally more interested and motivated to watch the animated movie. This was further confirmed by the extended duration for which children were able to maintain their attention to the movie (i.e. *∼*54 minutes) compared to the stories (i.e. *∼*36 minutes). Consequently, we were able to include more measurement conditions. To summarise, the use of an animated, age-appropriate movie could potentially result in improved response SNRs in hearing-impaired children. This, in turn, could enhance the reliability of neural tracking measures and reduce measurement time, both of which are crucial factors for clinical applications. Furthermore, the use of movies can aid children in maintaining attention, thus facilitating the inclusion of multiple measurement conditions within a single test session.

Although the magnitude of neural tracking might inherently vary between different stimuli, the overall results from both experiments were similar. This indicates that both stimuli can be suitable for obtaining an objective measure of speech intelligibility, as long as we examine relative differences within a subject, using the same speech material. Therefore, differences in reconstruction accuracy can be utilized to infer changes in speech intelligibility or the benefit of using a hearing aid. For example, significant changes in reconstruction accuracy between unaided and aided conditions may be a clear indication of improved speech intelligibility that may occur with the use of a hearing aid. Neural envelope tracking is shown to be quite robust, therefore being a good candidate for use in a clinical setting (for a review see: Brodbeck and Simon, 2020). However, measurement of neural tracking is a versatile tool as a wide range of speech processing abilities, including but not limited to envelope tracking, may be assessed in parallel (for a review see: Gillis et al., 2022b). For example, neural tracking of linguistic speech features has been suggested to provide a more accurate objective measure of speech intelligibility (Brodbeck and Simon, 2020; Gillis et al., 2021; Verschueren et al., 2022) and could be used to assess auditory as well as language abilities (Gillis et al., 2022b). However, as the neural processes targeted by these higher-order linguistic features are small, and the TRFs show a high between- and intra-subject variability, it is still challenging to extract a subject-specific measure. Further research is needed to explore the application of linguistic features in clinical practice.

In addition to selecting the appropriate speech stimulus and its features, the choice of analysis method (such as backward or forward modeling) should also be tailored to the intended purpose. This study revealed intriguing insights: reconstruction accuracies and TRF amplitudes remained unaffected by stimulus intensity in the aided condition. However, the latency of TRFs increased as stimulus intensity decreased, suggesting that TRFs could add valuable information about speech processing efficiency (Gillis et al., 2022a). On the other hand, decoders might be better suited for clinical applications, as backward models are fast, highly robust and do not require any channel selection or peak-picking methods.

In summary, our current findings highlight the potential of neural envelope tracking as a valuable tool for evaluating the speech perception abilities and hearing aid efficacy in younger individuals with hearing impairments. Furthermore, the use of narrated stories or engaging movies adds an element of enjoyment and familiarity, increasing the feasibility of implementing these measures in a clinical setting.

### Limitations

In our experiments, amplification was provided by children’s personal hearing aids. It is important to note that the functional status of the hearing aids at the time of testing was not analyzed either by electroacoustic analyses or real ear measurements. Therefore, we do not have an objective assessment of the gain or frequency response characteristics to confirm, at least in part, that children’s hearing aids were functioning properly or being optimally fitted at the time of testing. In addition, potential subtle differences in hearing aid circuitry and output might have contributed to the between-subject variability. However, since our primary focus is on comparing differences between conditions within the same individual, which is crucial for clinical implementation, we consider these potential differences to be relatively insignificant.

Another caveat of this study is that we only included a limited number of children in each experiment (N=8), which might restrict the ability to detect subtle or rare effects, particularly for the TRF analysis. As a result, caution must be exercised when interpreting the results, and further research with a larger and more diverse sample is recommended to further validate and strengthen the study’s conclusions. Nonetheless, the fact that the results were replicated in two separate experiments supports their validity. Hence, we are confident that the overall conclusions of this study remain unaffected by the small sample size.

## Conclusion

This research explored the potential of neural envelope tracking as an indicator of hearing function and hearing aid benefit in children with permanent hearing loss. The findings revealed several key insights. Firstly, we demonstrated that neural envelope tracking increased with increasing stimulus intensity in the unaided condition. However, in the aided condition, neural tracking remained stable across a wide range of stimulus intensities, as long as speech intelligibility was maintained. This was further supported by the TRF analysis, which showed that peak amplitude increased with stimulus intensity only in the unaided condition. Furthermore, our results provide the first evidence that personal hearing aids significantly improve neural envelope tracking, particularly in challenging speech conditions. Lastly, a strong correlation was observed between neural envelope tracking and behaviorally measured speech intelligibility. Altogether, our current findings highlight the potential of neural envelope tracking as a valuable tool for evaluating the speech perception abilities and assessing the effectiveness of hearing aids in younger individuals with hearing impairments. Furthermore, the use of narrated stories or engaging movies in the study design adds an element of enjoyment and familiarity, demonstrating the feasibility of implementing these measures in a clinical setting.

## Author statement

**Tilde Van Hirtum**: Formal analysis, investigation, data curation, writing-original draft, visualization and project administration. **Ben Somers**: Investigation, writing-review and editing, software. **Benjamin Dieudonné**: Writing-review and editing, software. **Eline Verschueren**: Investigation, writing-review and editing and project administration. **Jan Wouters**: Writing-review and editing, supervision and funding acquisition **Tom Francart**: Conceptualization, methodology, writing-review and editing, supervision and funding acquisition.

## Acknowledgements

The authors are grateful to all the children and their parents who made time to participate in this study. Our special thanks goes to the Centers for Ambulatory Rehabilitation in Flanders for their help with recruiting children, and to Jard Hendrickx and Ana Carbajal Chavez for their help in data acquisition. The authors would also like to thank Femke Vandenbempt (narrated stories) and Jeugdfilm Vlaanderen (animated movie) for providing the stimuli used in this study. Financial support was provided by the European Research Council (ERC) under the European Union’s Horizon 2020 research and innovation programme [grant number 637424, Tom Francart]; the KU Leuven research fund [grant number C3/20/045] and VLAIO innovation mandate (IM) [grant number HBC.2021.0203, Ben Somers].

