## Supplementary Information for "Neural envelope tracking predicts speech intelligibility and hearing aid benefit in children with hearing loss"

### Individual's behavioral speech intelligibility as a function of stimulus intensity for Experiment 1

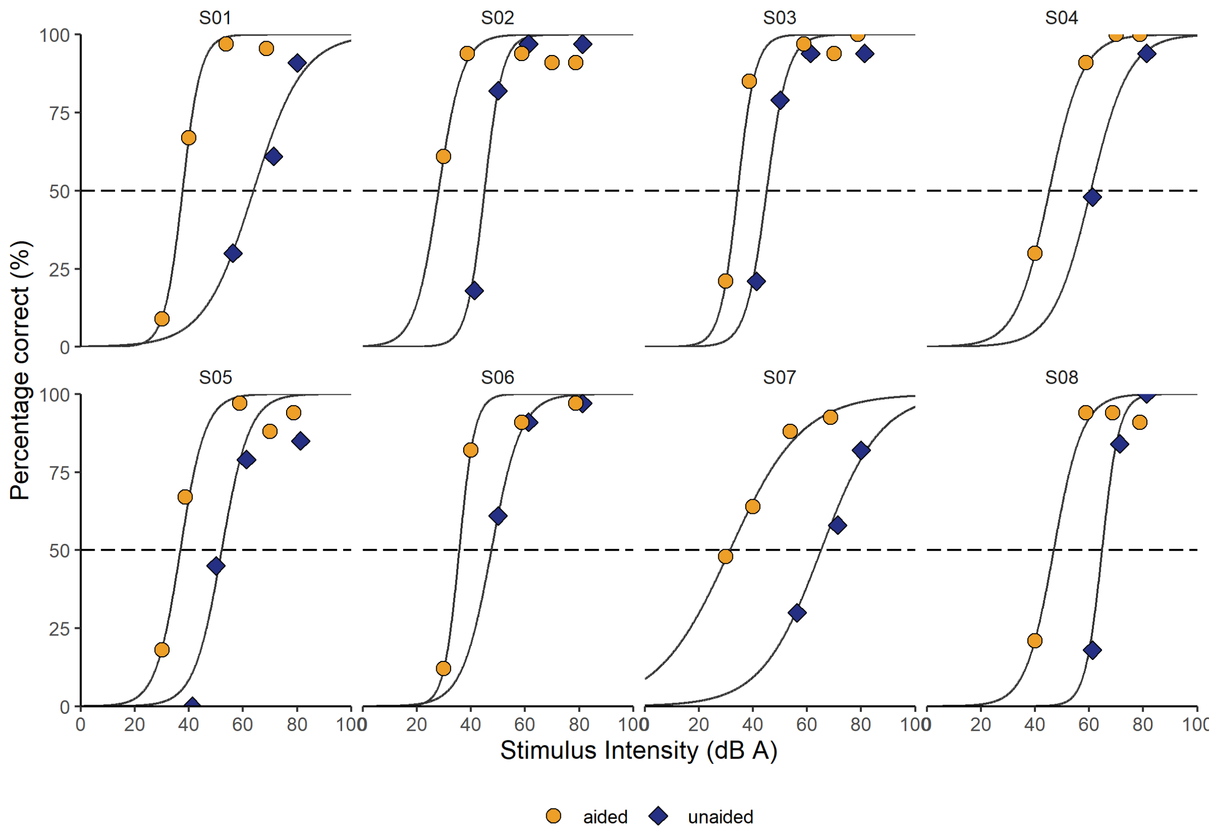


*Figure S1 Behavioral speech intelligibility of experiment 1. The individual's intelligibility (percentage phonemes correct) as a function of stimulus intensity for the unaided (blue diamonds) and aided (orange circles) condition, and the corresponding fitted performance intensity function from which the speech reception threshold (SRT) is derived.*

### Results of the TRF analysis for Experiment 1

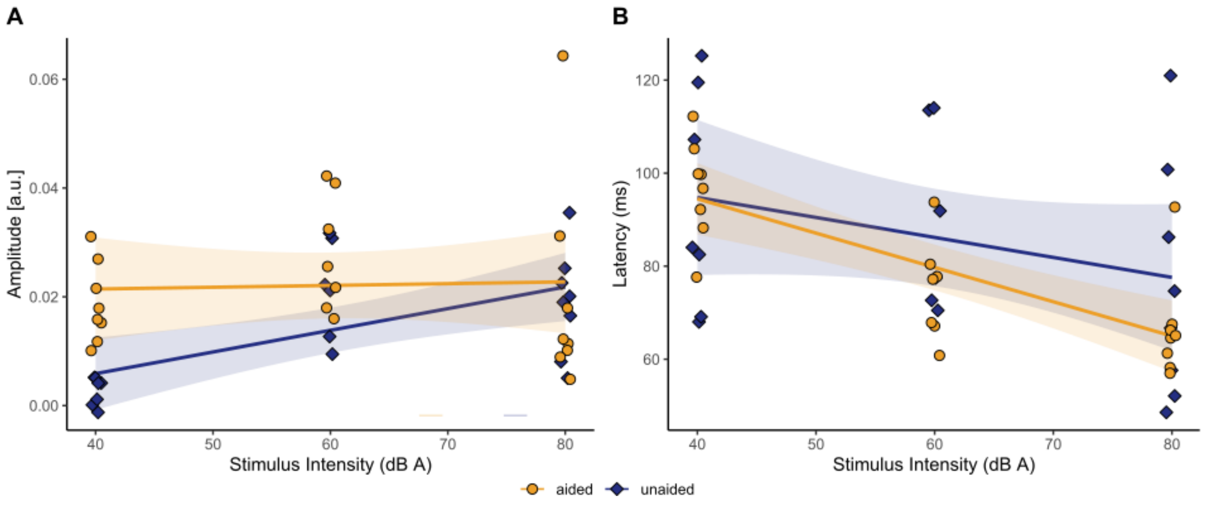


*Figure S2 Results of the correlation analyses of experiment 1. A. The effect of stimulus intensity on peak amplitude for the unaided (blue, diamonds) and aided (orange, circles) condition. Every datapoint represents an individual participant. The solid lines show the linear model fit. The shaded area represents the 95% confidence interval. B. The effect of stimulus intensity on peak latency for the unaided (blue, diamonds) and aided (orange, circles) condition.*

### Individual's behavioral speech intelligibility as a function of stimulus intensity for Experiment 2

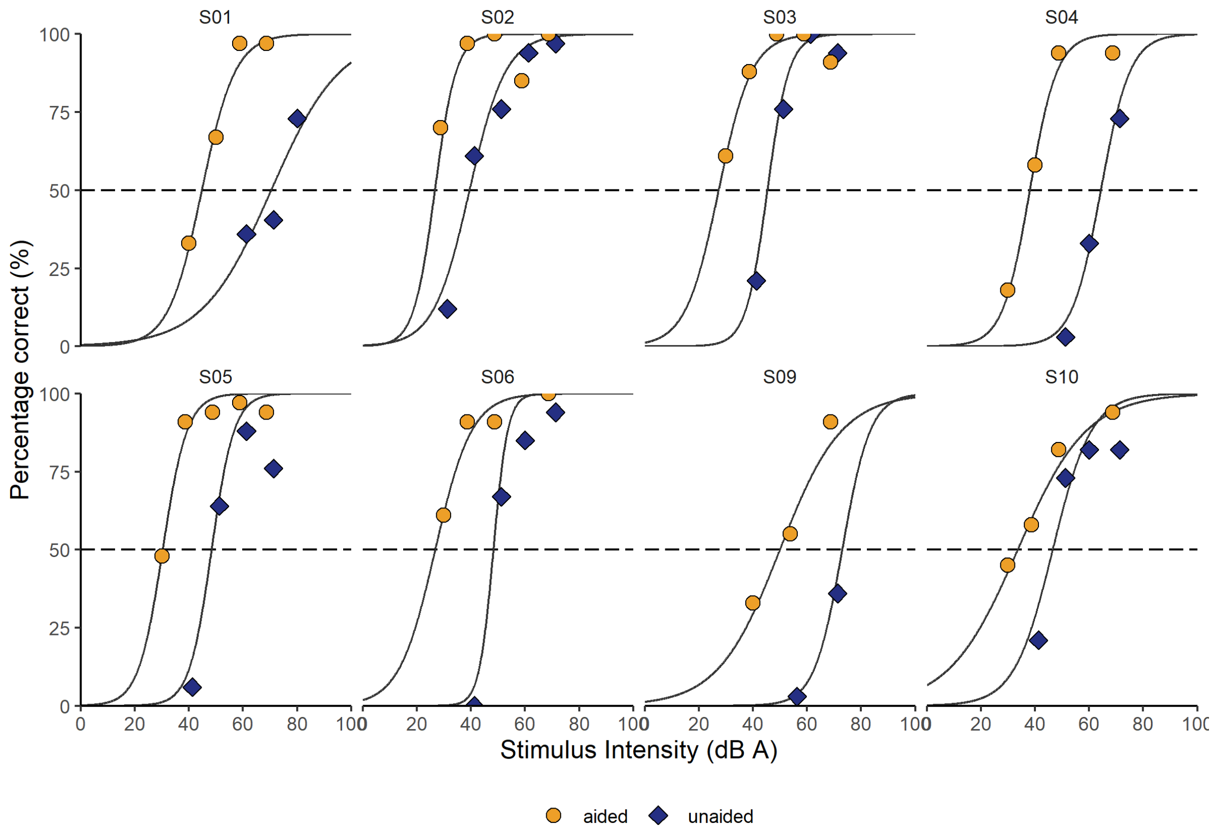


*Figure S3 Behavioral speech intelligibility of experiment 2. The individual's intelligibility (percentage phonemes correct) as a function of stimulus intensity for the unaided (blue diamonds) and aided (orange circles) condition, and the corresponding fitted performance intensity function from which the speech reception threshold (SRT) is derived.*

### Results of the TRF analysis for Experiment 2

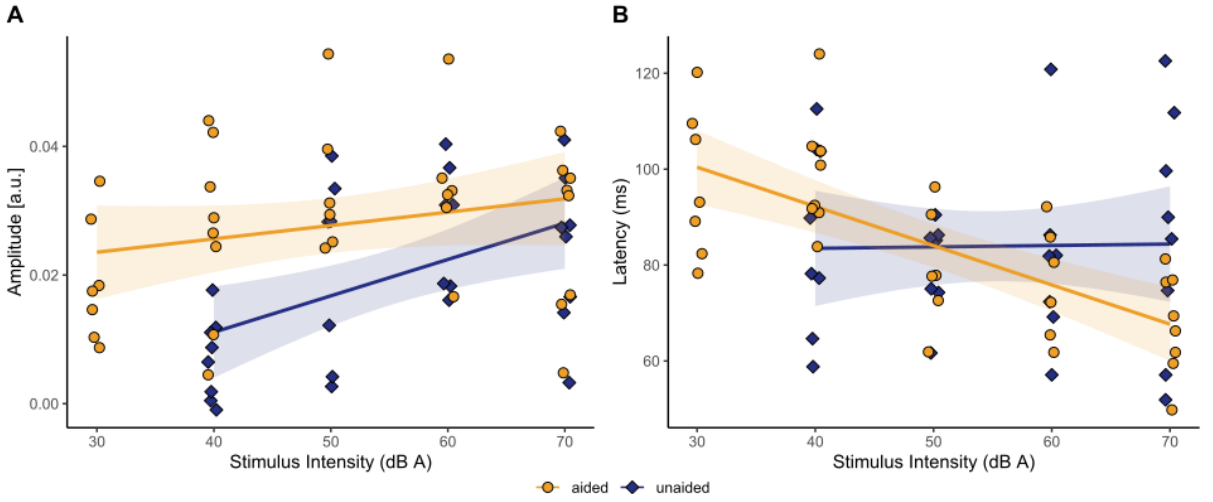


*Figure S2 Results of the correlation analyses of experiment 2. A. The effect of stimulus intensity on peak amplitude for the unaided (blue, diamonds) and aided (orange, circles) condition. Every datapoint represents an individual participant. The solid lines show the linear model fit. The shaded area represents the 95% confidence interval. B. The effect of stimulus intensity on peak latency for the unaided (blue, diamonds) and aided (orange, circles) condition.*

**Table S1 Output of LME model: effect of stimulus intensity and hearing aid use on envelope tracking in the delta band (Experiment 1)**

**Fixed-effect Terms**

| | **β-value** | **SE** | **df** | **t Ratio** | **p Value** |
|---|---|---|---|---|---|
| (Intercept) | 0.169 | 0.042 | 39 | 4.065 | <.001 |
| Intensity | 0.0002 | 0.001 | 39 | 0.317 | 0.7532 |
| Hearing aid | -0.300 | 0.057 | 39 | -5.278 | <.001 |
| Intensity:Hearing aid | 0.004 | 0.001 | 39 | 3.863 | <.001 |

**Table S2 Output of LME model: effect of stimulus intensity on TRF amplitude for unaided conditions (Experiment 1)**

**Fixed-effect Terms**

| | **β-value** | **SE** | **df** | **t Ratio** | **p Value** |
|---|---|---|---|---|---|
| (Intercept) | -0.01 | 0.0007 | 12 | -1.354 | 0.201 |
| Intensity | 0.0003 | 0.0001 | 12 | 3.389 | 0.005 |

**Table S3 Output of LME model: effect of stimulus intensity on TRF latency for unaided conditions (Experiment 1)**

**Fixed-effect Terms**

| | **β-value** | **SE** | **df** | **t Ratio** | **p Value** |
|---|---|---|---|---|---|
| (Intercept) | 101.582 | 19.453 | 12 | 5.222 | <.001 |
| Intensity | -0.204 | 0.308 | 12 | -0.664 | 0.519 |

**Table S4 Output of LME model: effect of stimulus intensity on TRF amplitude for aided conditions (Experiment 1)**

**Fixed-effect Terms**

| | **β-value** | **SE** | **df** | **t Ratio** | **p Value** |
|---|---|---|---|---|---|
| (Intercept) | 0.018 | 0.009 | 14 | 1.878 | 0.081 |
| Intensity | 0.0001 | 0.0002 | 14 | 0.489 | 0.632 |

**Table S5 Output of LME model: effect of stimulus intensity on TRF latency for aided conditions (Experiment 1)**

**Fixed-effect Terms**

| | β-value | SE | df | t Ratio | p Value |
|---|---|---|---|---|---|
| (Intercept) | 116.434 | 11.996 | 14 | -.706 | <.001 |
| Intensity | -0.565 | 0.189 | 14 | -2.995 | 0.009 |

**Table S6 Output of LME model: effect of stimulus intensity and hearing aid use on envelope tracking in the delta band (Experiment 2)**

**Fixed-effect Terms**

| | β-value | SE | df | t Ratio | p Value |
|---|---|---|---|---|---|
| (Intercept) | 0.089 | 0.035 | 57 | 2.544 | <.001 |
| Intensity | 0.002 | 0.001 | 57 | 3.730 | 0.7532 |
| Hearing aid | -0.296 | 0.058 | 57 | -5.113 | <.001 |
| Intensity:Hearing aid | 0.004 | 0.001 | 57 | 3.970 | <.001 |

**Table S7 Output of LME model: effect of stimulus intensity on TRF amplitude for unaided conditions (Experiment 2)**

**Fixed-effect Terms**

| | β-value | SE | df | t Ratio | p Value |
|---|---|---|---|---|---|
| (Intercept) | -0.015 | 0.008 | 21 | -1.834 | 0.081 |
| Intensity | 0.0006 | 0.0001 | 21 | 4.459 | <.001 |

**Table S8 Output of LME model: effect of stimulus intensity on TRF latency for unaided conditions (Experiment 2)**

**Fixed-effect Terms**

| | β-value | SE | df | t Ratio | p Value |
|---|---|---|---|---|---|
| (Intercept) | 75.387 | 18.499 | 21 | 4.075 | <.001 |
| Intensity | 0.179 | 0.329 | 21 | 0.544 | 0.592 |

**Table S9 Output of LME model: effect of stimulus intensity on TRF amplitude for aided conditions (Experiment 2)**

**Fixed-effect Terms**

| | β-value | SE | df | t Ratio | p Value |
|---|---|---|---|---|---|
| (Intercept) | 0.018 | 0.006 | 26 | 2.987 | 0.006 |
| Intensity | 0.0002 | 0.0001 | 26 | 1.586 | 0.125 |

**Table S10 Output of LME model: effect of stimulus intensity on TRF latency for aided conditions (Experiment 2)**

**Fixed-effect Terms**

| | β-value | SE | df | t Ratio | p Value |
|---|---|---|---|---|---|
| (Intercept) | 125.391 | 7.154 | 26 | 17.528 | <.001 |
| Intensity | -0.833 | 0.129 | 26 | -6.446 | <.001 |

**Table S11 Output of LME model: effect of stimulus choice on envelope tracking in the delta band (Comparison between Experiment 1 and 2)**

**Fixed-effect Terms**

| | β-value | SE | df | t Ratio | p Value |
|---|---|---|---|---|---|
| (Intercept) | 0.107 | 0.036 | 78 | 2.961 | 0.004 |
| Intensity | 0.002 | 0.001 | 78 | 3.554 | <.001 |
| Hearing aid | -0.274 | 0.049 | 78 | -5.597 | <.001 |
| Stimulus | 0.081 | 0.051 | 78 | 1.593 | 0.115 |
| Intensity:Hearing aid | 0.004 | 0.001 | 78 | 4.192 | <.001 |
| Intensity:Stimulus | -0.002 | 0.001 | 78 | -2.763 | 0.007 |
| Hearing aid:Stimulus | -0.041 | 0.025 | 78 | -1.658 | 0.101 |
